# ALMS1 contributes to centriole proximal architecture and stability

**DOI:** 10.64898/2026.07.15.738620

**Authors:** Sarah De Freitas, Maria Giovanna Riparbelli, Giuliano Callaini, Marine H. Laporte, Bénédicte Durand, Véronique Morel

## Abstract

Centrioles are highly organised microtubular scaffolds which grow and mature progressively during successive cell cycles. Their molecular organisation is extensively characterized, yet the contribution of several components to centriole assembly, maturation or stability is incompletely understood. Here, using ultrastructure expansion microscopy and transmission electron microscopy, we show that ALMS1, the protein mutated in Alström syndrome, is required for proper centriole architecture. In absence of ALMS1, RPE1 cells exhibit shorter centrioles with defects in the microtubular wall, including broken or missing triplets or open B/C tubules. These structural defects arise after procentriole assembly. We show that ALMS1 loss selectively reduces the proximal region proteins CCDC77 and CEP44, leaving intact central and distal ones. ALMS1 is further required for the recruitment of the proximal CEP135 cap and the clearance of the γ-tubulin/GCP2 pool present at the procentriole base. Our findings thus identify ALMS1 as a key organiser of the centriole proximal domain and required for remodelling and stabilising the proximal end of centrioles during cell cycle progression.

## Introduction

Centrioles are highly conserved microtubular organelles that come by pair and play key functions in cells. At the core of the microtubule organising centre, they are involved in cell division and organisation of the cell polarity. In non-dividing cells, the older of the two centrioles templates the cilium, which can be either non motile and involved in integrating signalling pathways, or motile and essential for cell movement and fluid displacement.

Centrioles present both proximo-distal and radial polarity. They are characterised by a nine-fold symmetry and are constituted, in mammals, by nine triplets of microtubules (A-B-C) which resolve into nine doublets towards the distal end of the centriole (Paintrand et al., 1992). This barrel of microtubules is decorated by numerous protein complexes, located internally, externally or aligned onto the microtubular wall and are key for the maintenance the centriolar structure and for its cellular functions (LeGuennec et al., 2021; Vasquez-Limeta and Loncarek, 2021). Alterations in centriole number or structure are associated with a variety of pathologies, including cancer and microcephaly, as well as a subset of ciliopathy-related conditions (Nigg and Holland, 2018).

During the G1 phase of the cell cycle, cells contain two related centrioles: a (older) mother centriole and a (younger) daughter centriole that was formed during the previous S-phase on the side of the mother centriole (Fig. 1A). Centriole duplication is tightly coupled to the cell cycle, resulting in the formation of two procentrioles per cell (Nigg and Holland, 2018; Gönczy, 2026). It is initiated at the onset of S-phase with the concentration of the kinase PLK4 as a single dot on the side of the pre-existing centrioles. This induces a well-orchestrated activation cascade, resulting in the formation of SAS-6 cartwheel. Its nine-fold symmetry serves as a blueprint for the nucleation of the procentriole microtubular wall (Dzhindzhev et al., 2010; Sonnen et al., 2013; Cunha-Ferreira et al., 2013; Moyer et al., 2015; Yamamoto and Kitagawa, 2019; McLamarrah et al., 2018; Dzhindzhev et al., 2014; Ohta et al., 2014; Kitagawa et al., 2011). Procentriole formation can be divided in two phases: the bloom phase during which both its length and diameter increase, and the elongation phase, which lasts until the end of the G2 phase. During this phase, its diameter is fixed and the procentriole only elongates (Laporte et al., 2024). Concomitantly to the growth of the microtubule triplets, several molecular scaffolds are formed that are involved in the formation or stabilisation of the centriole (Atorino et al., 2020; Pearson et al., 2009; Bayless et al., 2012; Schweizer et al., 2021; Bournonville et al., 2025). Among these, the pinhead contributes to the connection between the cartwheel and the microtubules and forms at the onset of their assembly; the A-C linker which marks the start of the elongation phase and bridges the A and C tubules of two neighbouring triplets of the proximal region of the centriole; and the inner scaffold which assembles during the late phase of procentriole formation and decorates the inner side of the centriole wall from the first proximal third to almost the distal end of the procentriole (LeGuennec et al., 2021). As the cell proceeds through G2 and M phase, the daughter centriole matures into a mother centriole and acquires distal appendages required for cilium formation. After mitosis, each daughter cell thus inherits a mature mother centriole associated to a newly formed daughter centriole.

**Figure 1:**
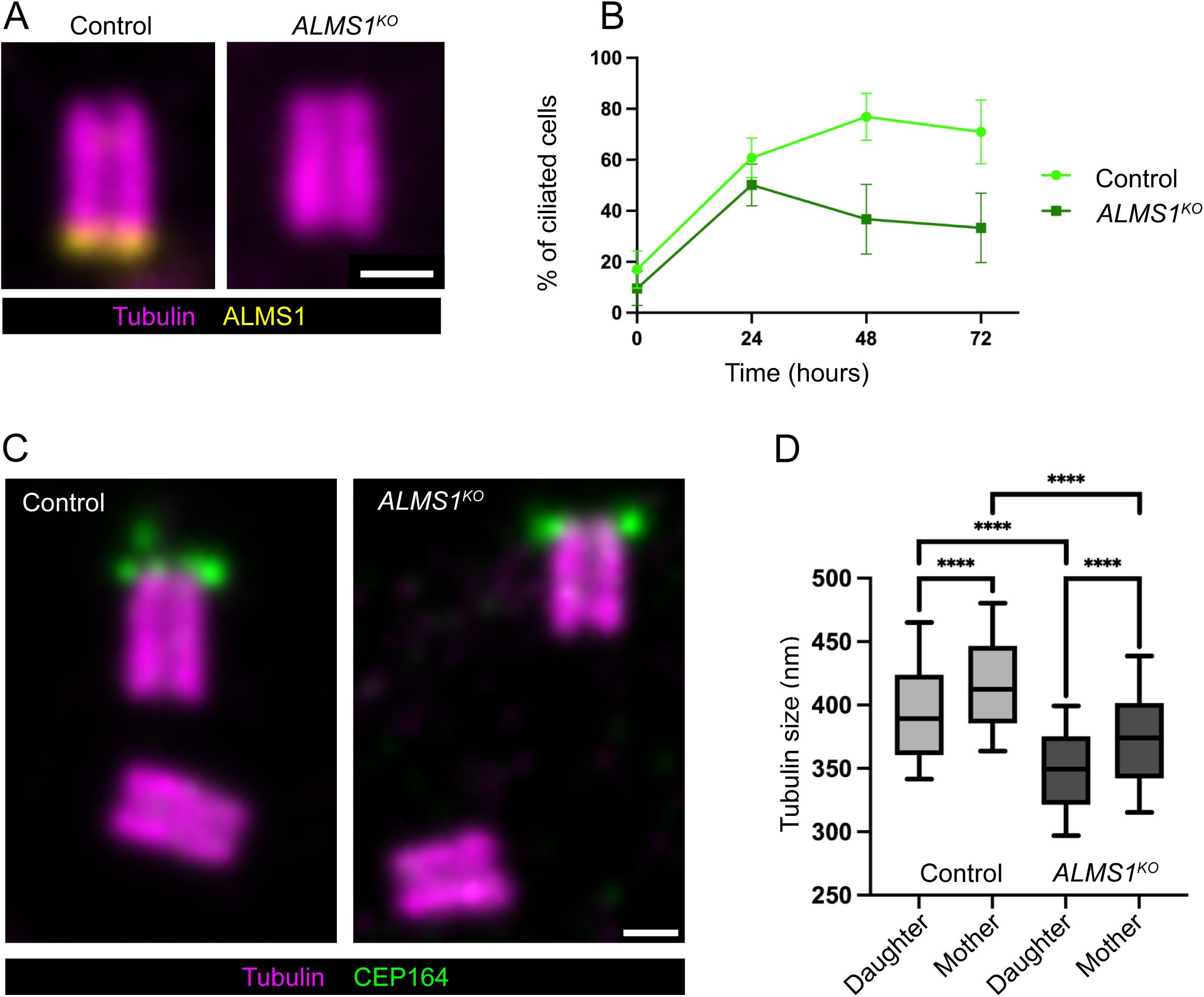
Loss of ALMS1 reduces the percentage of ciliated cells over time and centriole length. A- Expanded control and *ALMS1^KO^* centrioles in G1, shown in longitudinal view and stained for α/β- tubulin (magenta) and ALMS1 (yellow). B- Quantification of the percentage of ciliated cells over time following serum starvation (0h, 24h, 48h and 72h). In control cells (light green), the percentage of ciliated cells progressively increases over time and reaches a plateau of approximately 70% ciliation by 48 h. In ALMS*1^KO^* cells (dark green), the percentage of ciliated cells initially increases following serum starvation but from 24h onwards progressively decreases over time, reaching approximately 30% ciliation at 72 h. At all times the percentage of ciliated cells is lower in *ALMS1^KO^* than in control cells. C- Expanded control and *ALMS1^KO^* centrioles in G1, shown in longitudinal view and stained for α/β-tubulin (magenta) and CEP164 (green, labelling the distal appendages, specific of mother centrioles). D- Quantification of the tubulin length in G1. Daughter centrioles are ∼ 25 nm shorter than mother centrioles and *ALMS1^KO^* centrioles are ∼50 nm shorter than control centrioles (control daughter, n = 243; control mother, n = 226; *ALMS1^KO^* daughter, n = 276; *ALMS1^KO^* mother, n = 219). For all images, scale bars = 200 nm.

Among the proteins with a centriolar localisation, ALMS1 is a large protein of unknown structure besides a C-terminal ALMS domain involved in its localisation to the base of centrioles (Knorz et al., 2010). Mutations in *ALMS1* gene are responsible for the Alström syndrome, a rare ciliopathy associated with retinal degeneration, diabetes, obesity and dilated cardiomyopathy (Hearn, 2019; Hearn et al., 2005). We previously showed that *Drosophila* ALMS1 orthologs, Alms1a and Alms1b, are key players of centriole duplication in flies, involved in the activation loop between Plk4 and STIL/Ana2 required to trigger SAS-6 cartwheel assembly and subsequent procentriole formation. We further described ALMS1 localisation in RPE1 human cell line and established that it localises at the base of the procentriole from the elongation phase onwards (Brunet et al., 2025).

Here we took advantage of the *ALMS1^KO^* RPE1 cell line (Fig. 1A, (Woerz et al., 2024) to investigate ALMS1 contribution to centriole homeostasis. Using ultra-structure expansion microscopy (U-ExM) and EM analysis we show that in G1 phase, *ALMS1^KO^*centrioles are significantly shorter than control ones and present alteration of their tubular wall together with microtubule triplet defects. Analysis of centriole molecular organisation reveals that ALMS1 loss impacts the proximal domain of centrioles and its remodelling. In particular we observed a reduced A-C linker domain as well as loss of CEP135 proximal capping domain and altered Gamma tubulin redistribution in G1 daughter centrioles. Together our results demonstrate that ALMS1 is required for the organisation and maintenance of the proximal domain of centrioles required for its normal growth and structural stability.

## Results

### Loss of ALMS1 impacts cilium stability in RPE1 cells and centriole size

When cells exit the cell cycle and enter a quiescence (G0), the mother centriole docks to the plasma membrane and initiates axoneme formation, giving rise to a primary cilium (Fig. S1A). Loss of ALMS1, whether resulting from siRNA-mediated depletion, the use of a mutant cell line, or a mouse model, has previously been associated with a decrease in the proportion of ciliated cells, although these studies did not investigate changes occurring over time (Knorz et al., 2010; Woerz et al., 2024; Li et al., 2007). To further characterise the ciliation phenotype associated with ALMS1 loss (Fig. 1A), we first aimed to quantify the percentage of ciliated cells over time following serum starvation-induced quiescence (24 h, 48 h, 72 h, and 96 h). In control cells, the percentage of ciliated cells progressively increases from 17% at 0 h (fed state) to 60% after 24 h serum starvation and reaches a plateau of approximately 70% at 72 h and 96 h (Fig. 1B). In absence of ALMS1, we observed a lower percentage of ciliated cells compared to control, with 10% of fed cells presenting a cilium and approximately 50% of ciliated cells after 24 h starvation. The percentage of ciliated cells subsequently declined in *ALMS1^KO^*cells at 48 h and 72 h, whereas it remained stable in control conditions (Fig. 1B). These observations suggest that the decreased percentage of ciliated cells in *ALMS1^KO^*could result from both a decreased ability of *ALMS1^KO^* cells to initiate ciliation but also to maintain a cilium.

Since ciliogenesis and cilium stability relies on centriole proximo-distal ultrastructure and the correct assembly of the associated molecular scaffold (Fig S1)(Yang et al., 2005; Graser et al., 2007; Pearson et al., 2009; Bayless et al., 2012; Tsai et al., 2019; Curinha et al., 2025), we next investigated to which extent centriole organisation is impacted by ALMS1 loss. Taking advantage of the resolution gain obtained using Ultrastructure Expansion Microscopy (U-ExM) (Le Guennec et al., 2020), we observed that, in G1, *ALMS1^KO^* centrioles are shorter, by approximately 50nm, than control centrioles, both at the daughter and the mother stages (discriminated thanks to the presence of distal appendages labelled with CEP164 specific to the mother centriole, Fig. 1C,D). Hence, loss of ALMS1 impairs both primary cilium formation and stability and centriole size.

### ALMS1 contributes to the structural integrity of centriolar microtubule triplets

Beyond the reduction in centriole length, expansion microscopy revealed the presence of broken centrioles in *ALMS1^KO^* conditions with breaks either impacting the distal (Fig. 2A,B) or the proximal (Fig. 2C) ends of the centrioles. These observations were confirmed by transmission electron microscopy which further revealed alterations in the structure of the microtubule wall of centrioles in both longitudinal (15/23) and cross-sections (10/17) of *ALMS1^KO^* centrioles. Longitudinal section analysis identified shorter centrioles (Fig 2D,E) as well as breaks in the centriolar wall not only at the centriole ends (Figure 2F,G) but also in the medial regions of the centrioles (Figure 2H,I). Cross-section analysis revealed more subtle defects affecting the organisation of microtubule triplets including disconnected adjacent triplets suggestive of A-C linker discontinuity (Figure 2K, arrowhead), loss of C-tubules (Figure 2L,M arrow), single A tubule (Figure 2M, arrow) or open B and C tubules (Figure 2N,O, arrowheads).

**Figure 2:**
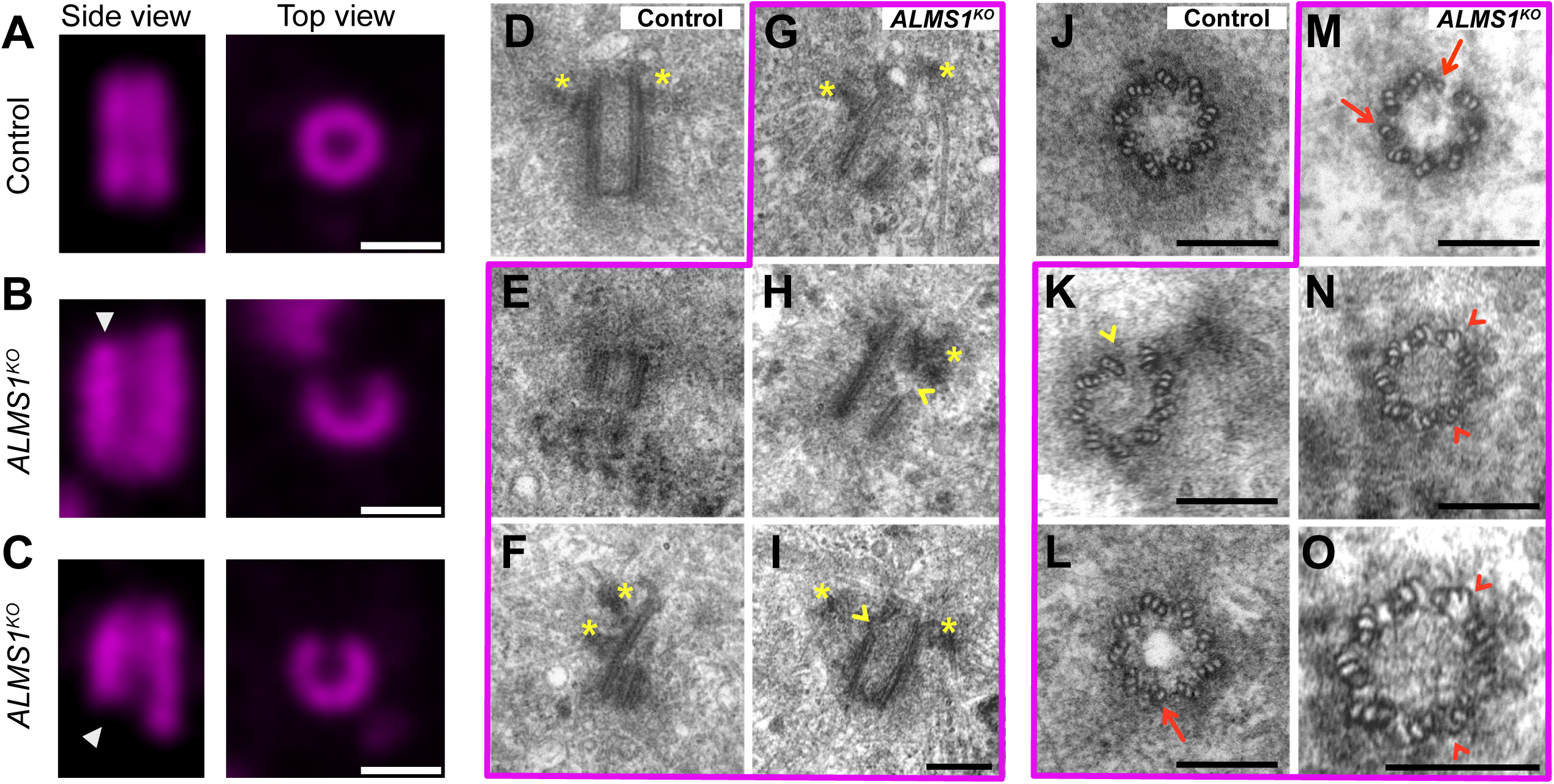
ALMS1 is required for the integrity of the centriole wall. A-C Expanded control (A) and *ALMS1^KO^*(B,C) centrioles in G1 stained for α/β-tubulin (magenta), shown in both longitudinal (side) and transverse (top) views. In *ALMS1^KO^* cells, discontinuities in the centriole wall are observed in both views and at both the distal (B, white arrowhead) and proximal (C, white arrowhead) ends of the centriole. D–P- Transmission electron microscopy (TEM) images of centrioles shown in longitudinal (D-I) and transverse (J-O) views. D,J, control centrioles. E-I and K-O, *ALMS1^KO^* centrioles. In *ALMS1^KO^*cells, defects in the centriole wall are observed at both the proximal and distal ends (arrowheads in F, I,K). In addition, the microtubule triplet architecture is altered, with some B- and C-tubules either missing (arrows in M and L) or incompletely closed (arrowheads N, O). O is a magnified view of N. asterisk: Distal appendages, in all images (U-ExM and TEM) scale bar = 200 nm.

Thus, ALMS1 appears to be required for the preservation of centriole architecture and the integrity of centriolar microtubule triplets.

### Procentriole formation is not affected by ALMS1 loss

ALMS1 forms a cap below the proximal end of centrioles (Fig. 1A, 3B) (Knorz et al., 2010; Brunet et al., 2025). This observation, together with the reduced G1 centriole size observed in *ALMS1^KO^* cells, led us to question whether centriole size defects might be a consequence of altered procentriole formation.

To address this, we followed procentriole growth in S and G2 using tubulin length as a first read-out of procentriole size. In addition, we followed the recruitment of SAS-6, the core component of the cartwheel, the pinhead protein CEP44, and the A-C linker protein CCDC77, previously shown to respectively precede, or localise simultaneously or after microtubule deposition (Fig. 3A)(Laporte et al., 2024). In both control and *ALMS1^KO^* conditions we observe a similar distribution of procentriole sizes, from 60 nm to 350 nm length (Fig. 3C). In agreement with the dynamics of localisation described by (Laporte et al., 2024) in U2OS cells, we observed a similar behaviour of SAS-6 during procentriole growth in control and *ALMS1^KO^* conditions, with SAS-6 domain growing until reaching a length of 150 nm and approximately 25 nm of SAS-6 sticking out of the procentriole proximally (Fig. 3D). As well, we do not observe any difference in the size of CEP44 or CCDC77 domains with respect to procentriole size in control and *ALMS1^KO^* cells (Fig. 3E,F). Note that, based on these quantifications, ALMS1 precedes A-C linker recruitment as it is first detected when the procentriole reaches a length of approximately 140 nm, while the A-C linker protein CCDC77 arrives on 180 nm long procentrioles (Fig. 3B,F). Hence, ALMS1 is recruited just before, or concomitantly to, the switch from the bloom to the elongation phase.

**Figure 3:**
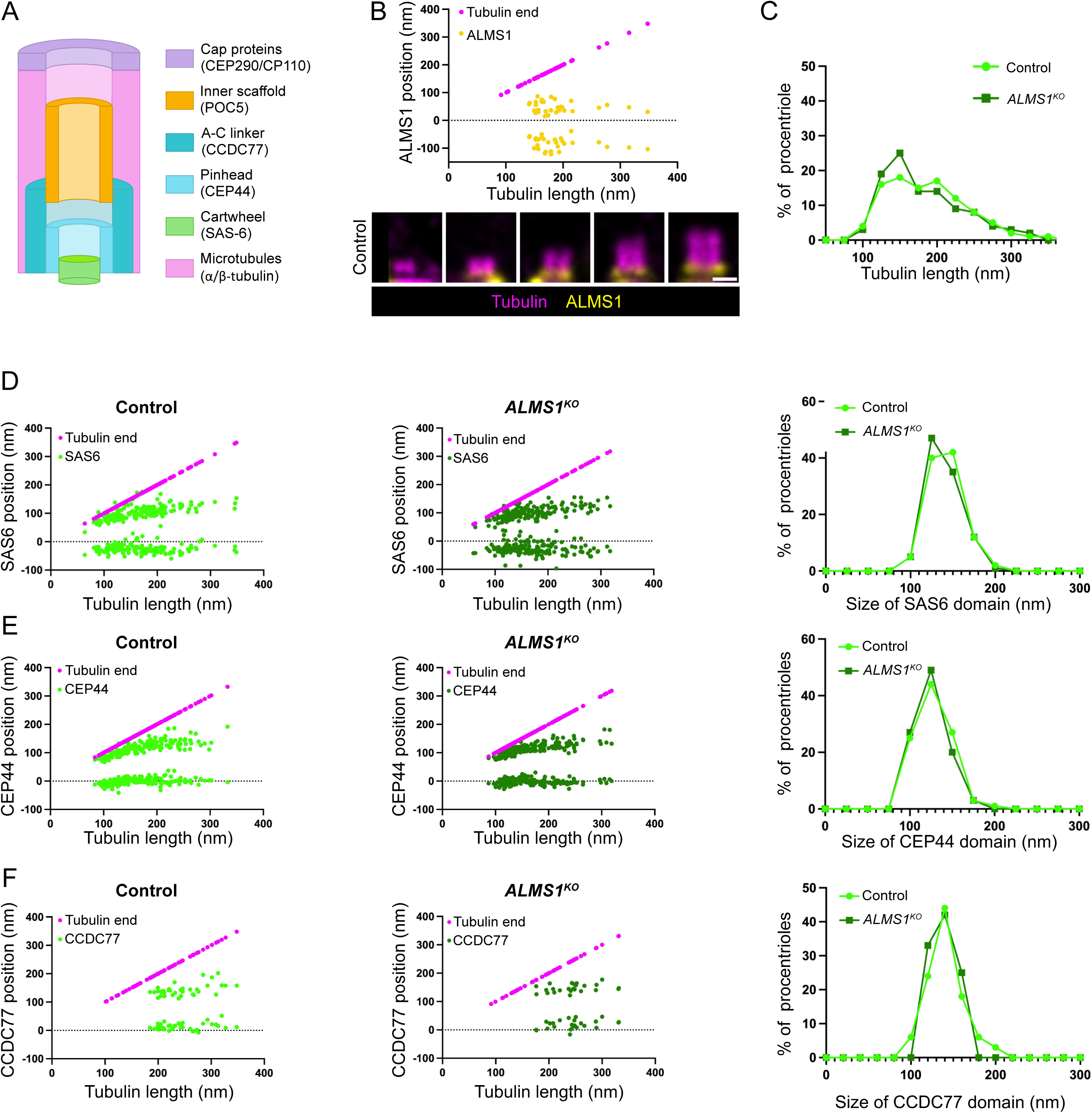
Procentriole growth and organisation in S phase is not affected in absence of ALMS1. A- Schematic representation of procentriole organisation in control cells during S/G2 phase with representative proteins used in this study. Cap proteins (purple, CP110 or CEP290), inner scaffold (dark orange, POC5), A-C linker (dark blue, CCDC77), pinhead (light blue, CEP44), cartwheel (SAS-6, light green), pericentriolar material (light orange) and microtubules (pink, α/β-tubulin). B- Longitudinal views of expanded control procentrioles labelled with α/β-tubulin (magenta) and showing the localisation of ALMS1 (yellow) during procentriole assembly in S phase. ALMS1 localises at the base of, and slightly overlaps with, the procentriole, starting as the procentriole reaches a length of 140 nm. Scale bars = 200 nm. C- Size distribution of procentrioles from control or *ALMS1^KO^* conditions. D-F- Progression of SAS6 (green, D), CEP44 (green, E) or CCDC77 (green, F) length relative to tubulin growth during procentriole assembly in control and *ALMS1^KO^* cell. The dashed line represents the procentriole proximal end, the magenta line the distal end of the centriole. The distribution of the centrioles according to SAS6, CEP44 or CCDC77 domain length is shown on the right. No significant differences were observed between control and *ALMS1^KO^* cells in the distribution of centrioles as a function of SAS6, CEP44, or CCDC77 domain length, indicating that the recruitment and localisation of these proteins during procentriole assembly are not affected by the loss of ALMS1.

Together, these observations thus suggest that procentriole growth proceeds normally in absence of ALMS1. As G1 centrioles are shorter in *ALMS1^KO^* than control, this further suggests that the size defects arise between the end of G2 and the beginning of G1 phase.

### ALMS1 is required for the integrity of the proximal centriole region

We next investigated whether subdomains of the centrioles are impacted upon ALMS1 loss when procentrioles have reached the G1 phase and become daughter centrioles.

The organisation of the distal end of the centriole, as reflected by CEP290 localisation, is not different between control and *ALMS1^KO^* conditions (Fig. S2). The inner scaffold domain (POC5) also has a similar size between control and *ALMS1^KO^* cells (Fig. 4A-A’). In contrast, the region proximal to POC5 is reduced in *ALMS1^KO^* cells (Fig. 4A’’). Both the pinhead (CEP44) and the A-C linker domains (CCDC77) are reduced in size in *ALMS1^KO^*condition compared to control (Fig. 4B-B’, 4C-C’). Last, CEP44 and CCDC77 domains extend below the proximal end of the daughter centriolar microtubules in *ALMS1^KO^* cells, a phenomenon not observed in control cells (Fig. 4B’’,C’’).

**Figure 4:**
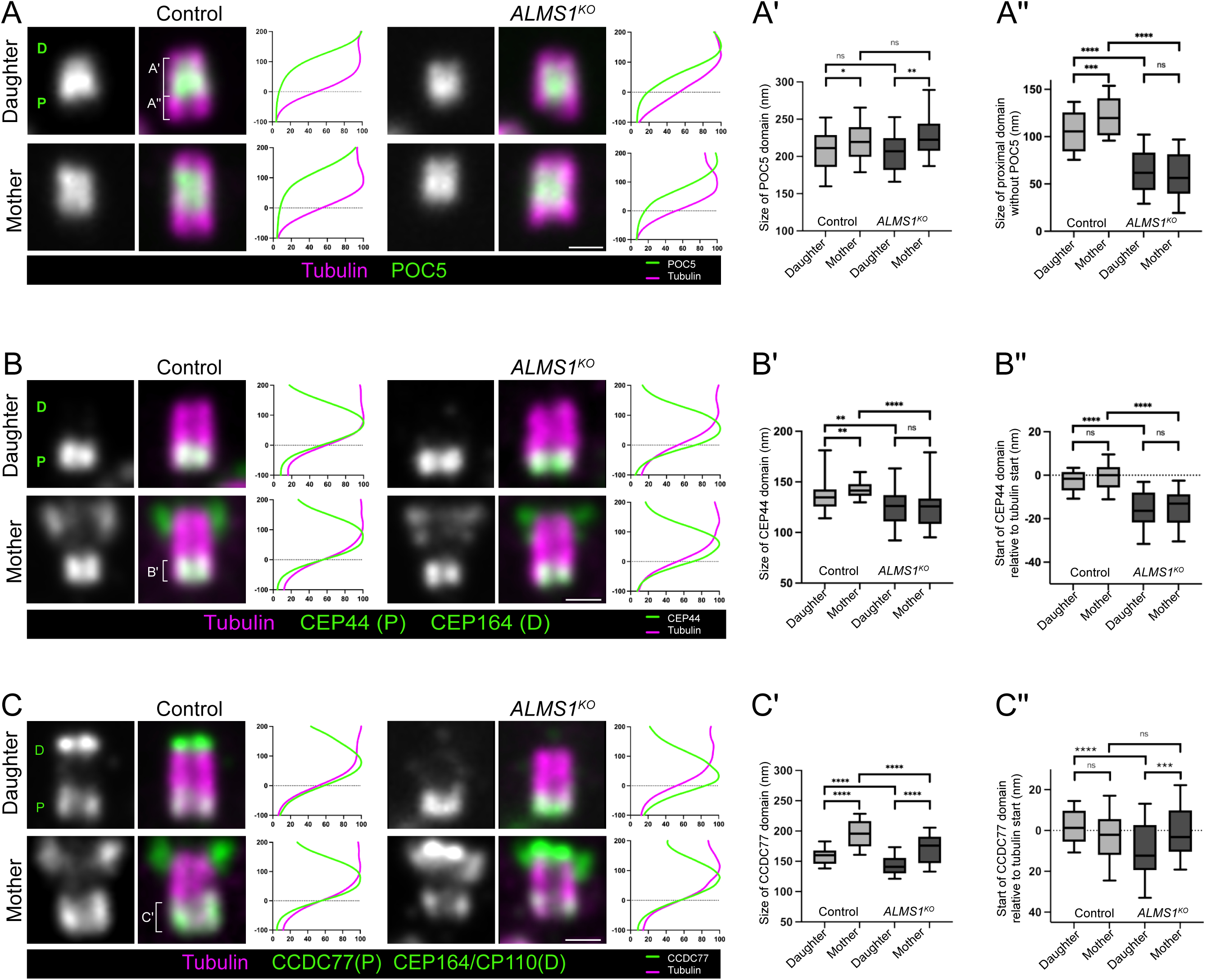
ALMS1 is required for the structural integrity of the proximal region of the centriole. A-C Longitudinal views of expanded G1 control and *ALMS1^KO^* daughter and mother centrioles labelled with α/β-tubulin (magenta) and showing the sub-centriolar localisation of POC5 (A), CEP44 (B) and CCDC77 (C) in green. CEP164 is used to identify the mother centrioles while CP110 marks the distal end of both daughter and mother centrioles. On the right, a plot profile of fluorescence intensity shows the position of POC5, CEP44 or CCDC77 relative to the proximal end of the centriole. A- The size of POC5 (green) domain and of the domain lying proximal to POC5 are depicted with a bracket and quantified in A’ and A”. A’- Box plot showing POC5 domain length A”- Box plot showing the size of the domain proximal to POC5 region (control daughter, n = 63; control mother, n = 62; *ALMS1^KO^* daughter, n = 65; *ALMS1^KO^*mother, n = 42). B- CEP44 (green) localises to the proximal part of the centriole and appears to occupy a smaller domain in *ALMS1^KO^* cells. B’- Quantification of CEP44 domain length. B”- Position of the start of CEP44 relative to the centriole proximal end. While CEP44 domains aligns proximally with the centriole in control daughter and mother centrioles, it is shifted below the centriole in absence of ALMS1. (control daughter, n = 46; control mother, n = 51; *ALMS1^KO^* daughter, n = 63; *ALMS1^KO^* mother, n = 43). C- CCDC77 (green) localises in the proximal part of the centriole and appears to occupy a smaller domain in *ALMS1^KO^* cells. C’- Quantification of CCDC77 domain length. The CCDC77 domain increases in length from daughter to mother centrioles in both control and *ALMS1^KO^* cells. It is significantly reduced in both daughter and mother centrioles of *ALMS1^KO^*cells compared with their respective controls. C”- Position of the start of CCDC77 relative to the centriole proximal end. While CCDC77 domains aligns proximally with the centriole in control daughter and mother centrioles, it is shifted below the centriole in absence of ALMS1 in daughter centrioles but aligns with the proximal end in *ALMS1^KO^* mother centrioles. (control daughter, n = 48; control mother, n = 48; *ALMS1^KO^* daughter, n = 38; *ALMS1^KO^* mother, n = 48). Scale bar: 200nm.

Hence, in absence of ALMS1 the medio-distal region of the daughter centriole is similar in size and organisation between control and *ALMS1^KO^* centrioles while the proximal pinhead and A-C linker domains are reduced, suggesting that the global shortening of *ALMS1^KO^* centrioles is sustained by a specific decrease in size of the proximal region. Together with the presence of a CEP44/CCDC77 domain devoid of tubulin proximally, this hints for a role of ALMS1 in the stabilisation of the centriole microtubules proximally.

### Loss of ALMS1 impacts proximal domain growth during daughter to mother centriole maturation

Observations by Chrétien et al. (1997) of purified centrosomes from KE67 cells suggest that centriole growth continues in G1 phase. Here we confirmed this hypothesis, as G1 mother centrioles are significantly longer than G1 daughter centrioles (by approximately 25 nm) both in control and *ALMS1^KO^* conditions (Fig. 1D). This prompted us to characterise the different centriole subdomains to determine whether they evolve during daughter to mother maturation and if their organisation is further altered in the absence of ALMS1.

In wild-type cells, we observed that the inner scaffold (labelled by POC5) and the pinhead proximal domain (labelled by CEP44) increase in size between the daughter and mother centrioles in G1 phase proportionally to that of the tubulin (Fig. 4A-B, S3A-B). However, the increase in size of the AC-Linker is proportionally greater than that of the tubulin between daughter and mother centriole (Fig. 4C, S3C). This suggests that during daughter to mother centriole growth, sub-centriolar domains growth is differentially controlled. These observations further suggest that the overlap between the A-C linker and the inner scaffold lengthen between daughter and mother centriole stages.

In absence of ALMS1, as in control conditions, mother centrioles are 25 nm longer than daughters (Fig. 1D). This growth comes with an increase of the POC5 domain in proportion to the centriole size and a relative larger increase of CCDC77 domain size (Fig. 4A-A’, C-C’, Fig. S3A,C). However, as opposed to control, this is not accompanied by an increase in the proximal region below POC5 (Fig. 4A’’). This absence of proximal domain extension in *ALMS1^KO^* cells, is also illustrated by the pinhead domain (CEP44) that does not grow in absence of ALMS1 compared to control between daughter and mother stages (Fig. 4B’). It also extends below the wall of both daughter and mother centrioles in *ALMS1^KO^* (Fig. 4B’’), whereas the A-C linker protein extends below the centriolar wall in daughter centrioles but aligns with microtubules proximally in mother centrioles (Fig.4C’’).

Together, these observations suggest an instability of the proximal end of the G1 daughter centriole in absence of ALMS1 that is present, and even worsens, from daughter to mother stages of the centriole. In addition, because the proximal domain (defined as proximal to POC5) increases in size in control but not in ALMS1 deficient centrioles, our observations also suggest that centrioles could grow from their proximal end during maturation from daughter to mother and that ALMS1 is required for this growth.

### ALMS1 is required for CEP135 recruitment as a proximal cap

Given that the proximal centriole region appears particularly affected in the absence of ALMS1, we investigated whether centriolar proximal protein CEP135 (Laporte et al., 2024; Kim et al., 2008; Matsuura et al., 2004), which has been identified as a proximity interactor of ALMS1 (Gupta et al., 2015), is impacted by ALMS1 loss. CEP135 exhibits several localisations within the centriole (Fig. 5A)(Laporte et al., 2024). In addition to a punctate localisation at the distal end of the centriole, a major pool of CEP135 is detected lining the inner side of the proximal wall of the centriole from the earliest stage of procentriole formation, while a second important pool is located below the centriole, forming a proximal CEP135 cap, and only observed when centrioles have reached the G1 stage. Here we show that *ALMS1^KO^* triggers a specific loss of the proximal CEP135 cap, while the distal and inner wall CEP135 localisations are not affected (Fig. 5A). This observation indicates that the absence of ALMS1 specifically disrupts the establishment or maintenance of the proximal CEP135 cap during G1 phase. These findings further support the hypothesis that ALMS1 plays a key role in maintaining the structure of the proximal centriolar region.

**Figure 5:**
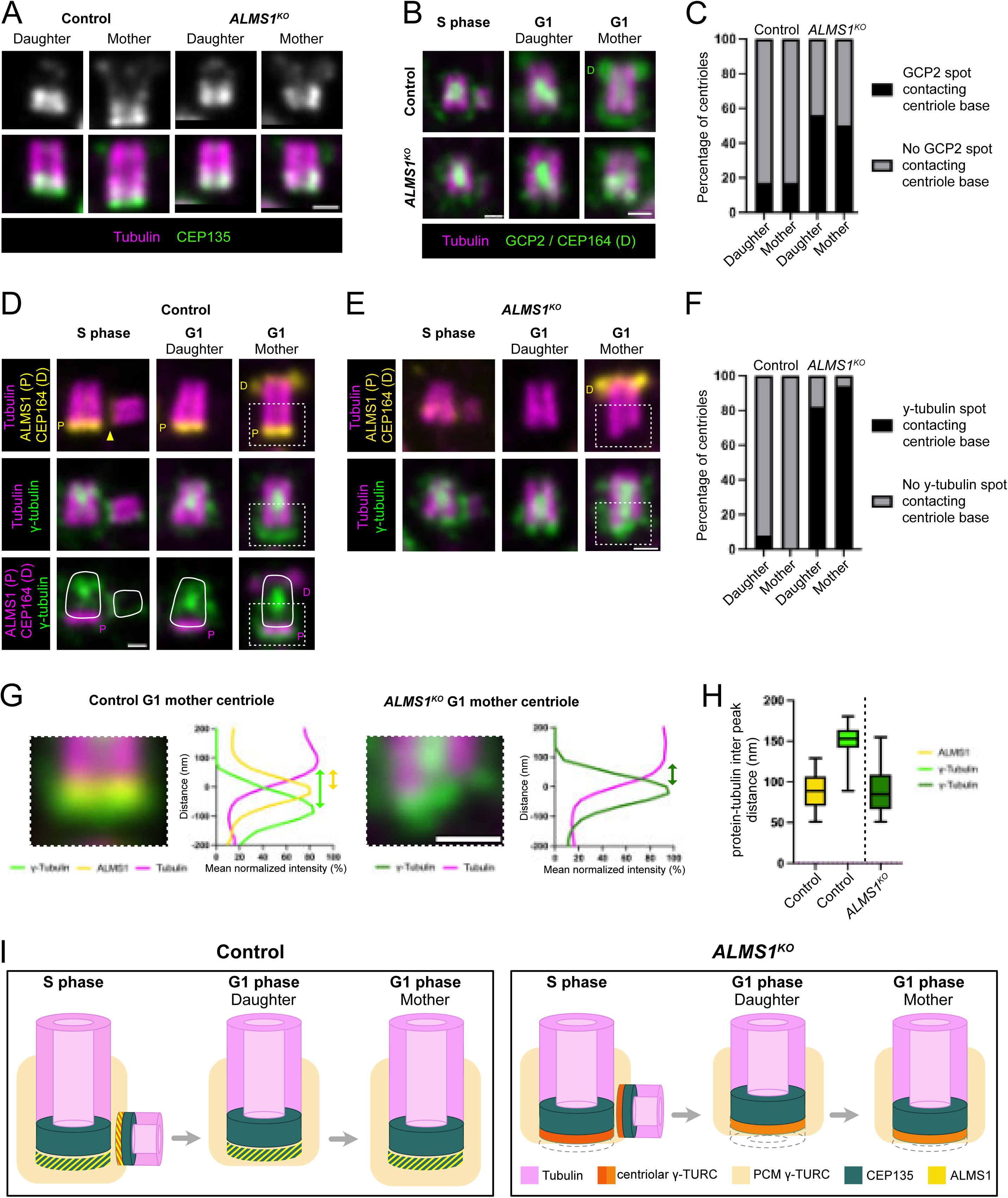
ALMS1 is required for the recruitment of CEP135 below the centriole and the removal of γ-tubulin during the S/G1 transition. A- Expanded centrioles in G1 stained for α/β-tubulin (magenta) and CEP135 (green). CEP135 localises at the proximal region of the centriole and extends into a platform-like structure below the centriole. In *ALMS1^KO^* cells, CEP135 localisation in the centriolar wall from the proximal region of the centriole is maintained but the proximal platform-like CEP135 signal is lost. B- Expanded control and *ALMS1^KO^* centrioles in S or G1 phase stained for α/β-tubulin (magenta) and GCP2 (green). In control cells, GCP2 displays multiple localisations: one signal associated with the inner scaffold of daughter and mother centrioles, a second PCM-like surrounding daughter and mother centrioles and a third signal at the base of the newly formed procentriole. In *ALMS1^KO^* cells, an additional GCP2 signal is detected below daughter and mother centrioles. C- Percentage of centrioles with GCP2 contacting the centriolar wall. D,E- Expanded control and *ALMS1^KO^* centrioles in S or G1 phase stained for α/β-tubulin (magenta), ALMS1 (yellow) and γ-tubulin (green). D- In control cells γ-tubulin localises as GCP2 in S and G1 phase. E- In *ALMS1^KO^* cells, an additional γ-tubulin signal is detected below daughter and mother centrioles. F- Percentage of centrioles with γ-tubulin contacting the centriolar wall. G- Mean plot profiles showing the relative localisation of γ-tubulin (green), ALMS1 (yellow) and tubulin (magenta) along the proximo-distal axis of the centriole (only the proximal half is shown here). In control centrioles, the γ-tubulin intensity peaks at approximately −80 nm, whereas the ALMS1 intensity peaks at approximately −10 nm. In *ALMS1^KO^* cells, the γ-tubulin intensity peak is shifted to approximately −10 nm. The Reference “0” corresponds to the 50% of tubulin intensity proximal pic value, used throughout thus work to define signal lengths. H- Box plot of distance between tubulin signal and ALMS1 or γ-tubulin, as measured by the distance between maximum intensity pics. The position of the tubulin pic is depicted as a magenta dashed line. I- Schematic representation of CEP135 and γ-tubulin localisation in mother and daughter centrioles in control and *ALMS1^KO^* cells. The diagram illustrates the loss of the proximal CEP135 platform in *ALMS1^KO^* centrioles while preserving CEP135 localisation along the centriole wall. It also depicts the canonical γ-tubulin localisation at the inner scaffold and PCM in control cells, together with the additional ectopic γ-tubulin accumulation below the mature centriole observed in *ALMS1^KO^*cells.

### ALMS1 is required for the removal of γ-Tubulin before G1 phase

The γ-TuRC (γ-tubulin ring complex) is the major complex responsible for nucleating the A-microtubule, while the B- and C-tubules are thought to form on the side of respectively the A- and the B-tubules, independently of the γ-TuRC complex (Guichard et al., 2010; Gupta et al., 2020). γ-TuRC complex is composed of numerous proteins, including 13 γ-tubulin and γ-tubulin complex proteins (GCP) (Guichard et al., 2010; Gupta et al., 2020; Tovey and Conduit, 2018; Zupa et al., 2021). In agreement with γ-tubulin localisations previously described in U2OS cells (Laporte et al., 2024), we observe that the yTURC proteins γ-tubulin and GCP2 exhibit a PCM-like localisation surrounding daughter and mother centrioles throughout the cell cycle (Fig. 5B,D), but also concentrate as a cap beneath the proximal end of the procentriole microtubules in S phase.

This concentration is not observed in daughter and mother centrioles present a pool of y-tubulin in the medio-distal part of the centriolar lumen (Fig. 5B-F) and a strong PCM localisation that surrounds the base and lateral proximal region of the centriole. Note that there is a gap between the centriolar γ-tubulin or GCP2 proximal pool and the microtubular wall that corresponds to the position of ALMS1 (Fig. 5G,H) in control cells.

In *ALMS1^KO^* cells, γ-tubulin and GCP2 are correctly recruited to the base of the procentriole during its formation, indicating that the early steps of γ-TuRC assembly are not affected by the absence of ALMS1 (Fig. 5B,E). However in *ALMS1^KO^* G1 cells, the proximal γ-tubulin/GCP2 pool below the centriole occupies (Fig. 5B,E,G) the position normally occupied by ALMS1 (Fig. 5G,H). This phenotype is observed in approximately 50% of centrosomes for GCP2 (Fig. 5C) and in more than 80% for γ-tubulin (Fig. 5F), suggesting that ALMS1 is involved in the rearrangement of the γTuRC complex from the procentriole to the daughter centriole stage. Note that neither the localisations of γ-tubulin/GCP2 associated with lateral PCM nor inside the centriolar lumen appear to be significantly affected by ALMS1 loss.

Together, these results indicate that ALMS1 plays a critical role in γ-TuRC rearrangement from the base of the procentriole before G1 phase (Fig. 5I).

## Discussion

Here, we demonstrate that ALMS1 loss affects the molecular structure of the centriole, resulting in a smaller footprint of its proximal domains. This includes a reduction in the A-C linker and pinhead domains, while the inner scaffold medio-distal domain remains unaffected. This is associated with a reduction in the size of the mother and daughter centrioles, as well as breaks in the centriolar wall and alterations of the microtubule triplets including B- and C- tubules loss, together suggestive of A-C linker defects. The precise timing of the observed molecular defects further suggests that ALMS1 contributes to the organisation of the centriole proximal end, and may either stabilise the centriole or promote its proximal growth.

The molecular alterations in CEP44, CEP135, CCDC77 observed in *ALMS1^KO^*centrioles raise the question of their contribution to observed overall structural defects. Depletion of either CEP44, CEP135 or both A-C linker proteins CCDC77 and WRD67 lead to shorter centrioles and centriolar wall breaks associated with loss of microtubule triplets or triplet integrity (Atorino et al., 2020; Lin et al., 2013; Bournonville et al., 2025). Specifically, depletion of A-C linker protein CCDC77 was previously reported to induce a decrease of daughter centriole size, broken centrioles and a reduction of CEP44 domain while inner scaffold protein POC5 domain size remained unchanged (Bournonville et al., 2025). These phenotypes share many similarities with what is observed upon ALMS1 depletion, suggesting that the reduction and instability of CCDC77 domain in *ALMS1^KO^* could be responsible for the altered structural stability of the centriole, as well as the shortening of the CEP44 domain. However, *ALMS1^KO^* does not fully recapitulate all described phenotypes associated with CCDC77 depletion. For example, CCDC77 siRNA leads to altered centriole duplication in 50% of cells, whereas we did not detect centriole duplication defects in *ALMS1^KO^* cells. As well, in *ALMS1^KO^* cells, CEP135 centriolar wall localisation is maintained whereas it is lost upon CCDC77 siRNA depletion (Bournonville et al., 2025). These differences might result from differences in the severity of CCDC77 perturbations between the two conditions, with a severe depletion of the protein after siRNA induced depletion compared to a reduction of the size of the CCDC77 domain or of its stability at the proximal end of the centriole in *ALMS1^KO^*cells.

If the pinhead-A-C linker structure is involved in the stability of the microtubule triplets, impairment of C-tubule formation by depleting epsilon or delta tubulin also impacts the A-C linkers (Atorino et al., 2020). This interdependency raises the possibility that the altered microtubular structure in *ALMS1^KO^*could be the cause rather than the consequence of A-C linker defects. In support of this hypothesis, we observed that procentrioles grow normally with CEP135, CEP44 and CCDC77 being assembled similarly in *ALMS1^KO^* and control centrioles. The first differences between these two conditions are observed in G1 daughter centrioles where the proximal domain is reduced in *ALMS1^KO^* compared to control. We also observe a slight proximal extension of CCDC77 below the centriole, while it is normally always associated with microtubular wall. This lagging localisation is resolved in mother centrioles where CCDC77 aligns with the microtubular wall proximally. Together, these observations suggest that in absence of ALMS1, microtubules might be destabilised and partially disassembled proximally, which could in turn lead to a remodelling of the A-C linker.

In *ALMS1^KO^* cells, the CEP135 proximal cap, normally observed from the end of the procentriole stage, is absent, while it is still present in absence of CCDC77. This suggests that this proximal CEP135 domain does not depend on CCDC77 and could be directly under ALMS1 control. More, a γTURC pool is observed below the centriolar wall, a localisation characteristic of growing procentrioles that is lost in G1 daughter and mother centrioles. Thus, in the absence of ALMS1, the proximal cap retains an early procentriole-like organisation, which could reflect an “immature” state and be responsible for the proximal microtubule instability. It has been shown that the yTURC which caps the A-tubule of procentrioles is removed during their elongation (Guichard et al., 2010). This was also suggested by UExM observations of procentriole assembly (Laporte et al., 2024). Our observations during the maturation of procentrioles into daughter G1 centrioles also indicate that the initial proximal pool is replaced by an enlarged pool separated from the centriolar wall by ALMS1, likely corresponding to the PCM-associated yTURC. In cells depleted of ALMS1, we hypothesise that the yTURC cap on the procentriole is not removed during procentriole maturation. However, it is also possible that the PCM-associated yTURC has colonised the space left by the absence of CEP135 and ALMS1. While we cannot completely exclude this latter hypothesis, close examination of the proximal yTURC domain in ALMS1-depleted centrioles reveals that it does not completely surround the proximal centriole as PCM material does. In addition, our observations suggest that the removal of yTURC, as timely described in Laporte et al, and the recruitment of proximal CEP135 cap follow ALMS1 recruitment at the procentriole. It is therefore tempting to speculate that ALMS1 stabilises alone, or together with CEP135, the microtubule triplets during centriole life. Whether yTURC maintenance is directly controlled by ALMS1 or a consequence of defective recruitment/stabilisation of CEP135 proximal cap remains to be investigated.

It is commonly accepted that three cell cycles are necessary for a centriole to become fully mature (Sullenberger et al., 2020). Here, by precisely characterising the size and molecular organisation of daughter and mother centrioles in G1, we show that during the second cell cycle the daughter centriole, in addition to acquiring distal appendages, continues to grow and is subjected to a resizing of the footprint of several domains, with a larger increase of the A-C linkers domain compared to the Inner scaffold. Extrapolating from the size of each domain and their relative position to centriole proximal end, we conclude that the A-C linker and Inner scaffold overlap increases from 13% to 18 % between daughter and mother centriole, in agreement with the previously published overlap estimation of 19% for all mature centrioles (daughter or mother centrioles altogether, (Laporte et al., 2024). How this remodelling is orchestrated during centriole maturation is still enigmatic, but our work also reveals that ALMS1 is involved (either directly or indirectly) in the stabilisation of the microtubules as well as in the reorganisation of the domain below the centriole as illustrated by the maintenance of the yTURC concentration and the lack of CEP135 cap.

Last, in the literature, ALMS1 loss is associated with premature disjunction of mother and daughter centrioles in G1 and defective cilia formation, and, as shown here, altered cilia stability (Knorz et al., 2010; Li et al., 2007; Woerz et al., 2024). Notably, CEP135, which localisation is altered in *ALMS1^KO^* cells, is required for the recruitment of C-NAP1 at the proximal end of the centriole which in turn mediates the assembly of inter-centriolar fibres and the rootlet, respectively involved in daughter and mother centriole connection in G1 and cilium anchoring and maintenance (Bahe et al., 2005; Kim et al., 2008; Vlijm et al., 2018; Yang et al., 2005, 2006; Ryu et al., 2024; Theile et al., 2023). In absence of ALMS1, lack of CEP135/C-NAP1 recruitment at the base of the centriole could thus explain cilium instability, without excluding a contribution of altered structural integrity of *ALMS1^KO^* centrioles to defective cilia homeostasis.

Here we show that loss of ALMS1 leads to centriole size decrease and altered centriole microtubular structure. We further show that ALMS1 is involved in the reorganisation of the proximal domain of daughter centrioles, including the recruitment or stabilisation of the CEP135 proximal cap. We therefore propose that ALMS1 contributes to the organisation of the centriole proximal end, either by stabilising or by promoting its growth. Reaching a decision between these two options will require greater temporal and molecular resolution, but our work reveals ALMS1 to be a key factor in centriole maturation and integrity.

## Material and Methods

### Human cell culture

hTERT RPE1 Cas control and *ALMS1^KO^* (Human immortalized Retinal Pigment Epithelial cells; gift from T. Beyer (Woerz et al., 2024)) cells were grown on non-coated 12 mm diameter coverslips in DMEM-F12 Glutama× 10% FBS and penicillin/streptomycin 0,1% at 37°C in 5% CO2. RPE-1 were seeded at 200 000 cells/mL or 240 000 cells/mL in 6-well plates for 24 h before being processed for U-ExM or immunofluorescence.

RPE1 *ALMS1^KO^* cell line was generated by (Woerz et al., 2024) and carries two points mutations in ALMS1 exon 8 and 10 resulting in a premature stop of ALMS1 coding sequence and the absence of ALMS1 protein (Fig. 1A).

### Ciliogenesis quantification

24 hours after cell seeding, the culture medium was replaced with DMEM/F-12 GlutaMAX supplemented with 0.5% FBS and 0.1% penicillin/streptomycin to induce serum starvation, thereby promoting primary cilium formation. 24, 48, and 72 hours after serum starvation, cells were fixed in Methanol and processed for immunofluorescence as described below. The percentage of ciliated cells corresponds to the number of ciliated cells (as estimated with Arl13B labelling) over the total number of cells (as estimated with Hoechst labelling).

### Ultrastructure Expansion Microscopy (U-ExM)

U-ExM was performed as described previously described (Le Guennec et al., 2020), without pre-fixation step. Coverslips were incubated in 2% AA + 1.4% FA diluted in PBS for 3 to 5 h at 37°C prior to gelation in monomer solution (19% sodium acrylate, 0.1% bis-acrylamide, 10% acrylamide, 0.5% TEMED) supplemented with APS (final concentration 0.5%) for 30 min at 37°C. Gels were cut using a 0.4 mm biopsy punch prior denaturation in (SDS 200mM, NaCl 200mM, Tris pH9 50mM) for 1.5h at 95°C. Gels were washed 3×20 min in ddH_2_O. At this stage, gels were either stored at −20°C in 70% glycerol / 30% ddH₂O or directly processed for staining as follows.

### Antibody staining

Gels were rinsed in PBS (3 × 20 min) and incubated with primary antibodies in PBS-BSA 2% at room temperature overnight with agitation. After 3 × 20 min washes in PBS-0.1% Tween gels were incubated with secondary antibodies (1/1000) and Hoechst (1/500) in PBS-BSA 2% at room temperature overnight with agitation and washed 3 × 20 min in PBS-0.1% Tween prior expansion in ddH₂O (3 × 20 min).

For classical immunofluorescence, cells were fixed at −20°C in 100% methanol for 5 minutes, followed by 3×5min washes in PBS 1X. Permeabilisation in PBS-0.1% Tween (20min at RT) followed by a blocking step in PBS-2% BSA (20min at RT) was done before incubation 1h at RT in PBS-BSA2% with primary antibodies. Cells were then washed 3×5min in PBS-0.1% Tween, incubated 1h at RT in secondary antibodies and Hoechst diluted at 1/1000 in PBS-2% BSA, washed 3×5min in PBS-0.1% Tween at RT and mounted in Vectashield for imaging.

### Antibodies

See Table for details regarding primary antibodies. Secondary fluorescent antibodies were purchased from Invitrogen (A11008, A11029, A11076 and A21244), Hoechst H33342 (20mg/mL).

| ANTIBODIES |  |  |
| --- | --- | --- |
| GUINEA-PIG ANTI-BETA TUBULIN | ABCD antibodies | AA344 |
| GUINEA-PIG ANTI-ALPHA TUBULIN | ABCD antibodies | AA345 |
| RABBIT ANTI-POC5 | Proteintech | A303-341A |
| RABBIT ANTI-CCDC77 | Proteintech | 26369-1-AP |
| RABBIT ANTI-CEP44 | Proteintech | 24457-1-AP |
| RABBIT ANTI-CEP164 | Proteintech | 22227-1-AP |
| MOUSE IGG2A ANTI-CEP164 | Santa Cruz Biotechnology | sc-515403 |
| RABBIT ANTI-CEP290 | Proteintech | 22490-1-AP |
| RABBIT ANTI-SAS6 | Proteintech | 21377-1-AP |
| RABBIT ANTI-CEP135 | Proteintech | CL488-24428 |
| MOUSE IGG1 ANTI-GAMMA-TUBULIN | Sigma | GTU88 _ T5326 |
| RABBIT ANTI-ALMS1 | Proteintech | 27231-1-AP |
| RABBIT ANTI-GCP2 | Proteintech | 25856-1-AP |
| **ANTI ARL13** | Gift from C. Brun |  |
| **HOECHST** | Sigma | H33342 |

### Imaging

Expanded gels were mounted onto 24 mm coverslips coated with poly-D-lysine (0.1 mg/ml). For centriole substructure analysis, gels were imaged using an IX 83 inverted microscope from Olympus, equipped with a Yokagawa CSU-X1 Spinning Disk Unit, Borealis technology for homogeneous illumination and Ixon3 888 EM-CCD camera from Andor. The oil immersion Plan Apochromat 60x/1.42 NA objective from Olympus was used for all acquisitions. For ciliation analysis, cells were imaged using a Leica Thunder Imager microscope with an oil immersion HC Plan Apochromat 63x/1.40 objective.

### Transmission Electron microscopy (TEM)

*ALMS1^KO^* and control cells were plated on microscope glass coverslips and fixed overnight at 4°C in 2.5% [v/v] glutaraldehyde in 0.1M phosphate buffer (pH 7.2, PBS). After washing for 30 min in PBS, the cells were post-fixed in 1% osmium tetroxide in PBS for 1 h. The cells were then dehydrated in a graded series of ethanol, and infiltrated with a mixture of Epon–Araldite resin. After polymerization for 48 h at 60°C the coverslips were removed from the resin with a short immersion in liquid nitrogen. Ultrathin sections (50–60 nm thick) were cut with a LKB Ultratome NOVA, equipped with a diamond knife. The sections were collected with copper slot grids coated with formvar (1% in chloroform). After drying with filter paper, the sections were stained with 2% aqueous uranyl acetate for 20 min in the dark, and then with lead citrate for 2 min. The preparations were observed with a Philips CM10 electron microscope at 100 kV, equipped with a Morada CCD camera (Olympus, Tokyo, Japan).

### Quantification and statistical analysis

#### Longitudinal measurements from side viewed centrioles

Measurements of centriole length, relative positioning and protein coverage were performed on co-labeled images, with tubulin consistently serving as a reference marker to estimate centriole length. The ‘‘PickCentrioleDim’’ plugin (https://github.com/CentrioleLab) was used to assist in identifying the start and end points of the fluorescent signal, defined as 50% of the maximum intensity at each end of the centriole. This approach automatically generates a dataset containing the coordinates of signal boundaries for each analysed channel. For length values, the distance between the fluorescent signal extremities was calculated. All shown values are corrected for the gel expansion factor.

#### Calculation of the expansion factor

For each gel analysed, the mean nuclear area was measured: nuclei were labelled by Hoechst, imaged using the 10x objective of the Andor spinning disk microscope and their area was quantified using Fiji. For each gel, the expansion factor was calculated as the mean expanded nuclei area divided by the mean nuclei area from not expanded cells of the same genotype. More than 1000 nuclei were analysed to generate the not expanded reference. This approach was chosen because as centriole structure is altered in the mutant cell lines, it cannot serve as an internal reference for expansion measurements.

#### Image processing

Images shown were resized with Fiji to obtain a pixel size of 10nm to improve precision.

#### Statistical analysis

Wilcoxson exact tests were performed using R. All the graphs were generated on GraphPad Prism10.

| CHEMICALS | SUPLIERS | REFERENCES |
| --- | --- | --- |
| ACRYLAMIDE (AA, 40%) | Sigma | A4058 |
| FORMALDEHYDE (FA, 36.5–38%) | Sigma | F8775 |
| SODIUM ACRYLATE (SA, 97–99%) | Sigma | 408220 |
| N, N-METHYLENBISACRYLAMIDE (BIS, 2%) | Sigma | M1533 |
| TETRAMETHYLETHYLENDIAMINE (TEMED) | Invitrogene | 15524-010 |
| AMONIUM PERSULFATE (APS) | Euromedex | EU0009-A |
| SODIUM DODECYL SULFATE 20% (SDS) | Euromedex | EU0660 |
| METHANOL | Fisher Scientific | M/4056/15 |
| TWEEN-20 | Euromedex | 1005-7 |
| BOVIN SERUM ALBUMIN (BSA) | Sigma | P1379 |
| GLYCEROL | Euromedex | EU3550 |
| POLY-D-LYSINE | Sigma | P1024 |
| VECTASHIELD | VECTOR | H-1900 |
| DMEM/F12 | PAN BIOTECH | P04-41150 |
| FETAL BOVINE SERUM (FBS) | Dutscher | S1400-500A |
| PENICILLIN / STREPTOMYCIN | PANbiotech | P06-07100 |
| TRYPSIN-EDTA 0,25% | Gibco | 25200-056 |
| GLUTARALDEHYDE 25 % | Sigma-Aldrich | G5882 |

## Acknowledgments

This work was supported by the ANR CiliAS. SdF was supported by a doctoral fellowship from the FRM. ThE authors acknowledge the contribution of the SFR Santé Lyon-Est (UAR3453 CNRS, US7 Inserm, UCBL) facility: CIQLE (a LyMIC member). We thank T. Beyer, M. Ueffing and K. Boldt for generously sharing the ALMS1^KO^ and control hTERT RPE1 cell lines. We thank J. Thomas for her help with manuscript proof reading.

---

**Figure S1: Schematic representation of the cell cycle and centriole molecular organisation.**

A- Scheme illustrating the centriole biogenesis with respect to the cell cycle phases. B- Schematic representation of the S/G2 procentriole, G1 daughter and mother centrioles showing the sub-centriolar domains organisation with representative proteins used in this study. Cap proteins (purple, CP110 or CEP290), Distal appendages (CEP164, dark grey), Sub-distal appendages (light grey), inner scaffold (dark orange, POC5), A-C linker (dark blue, CCDC77), pinhead (light blue, CEP44), cartwheel (SAS-6, light green), pericentriolar material (light orange) and microtubules (pink, α/β-tubulin).

**Figure S2: Localisation of cap protein CEP290 in control and *ALMS1^KO^* G1 centrioles.**

Expanded centrioles in G1 stained for α/β-tubulin (magenta) and CEP290 (green). CEP290 caps the distal end of both daughter and mother centrioles in G1. No differences were observed between control and *ALMS1^KO^*conditions. Scale bar: 200nm.

**Figure S3: Quantification of POC5, CEP44 or CCDC77 domain length relative to centriole length.**

A- Percentage of centriole covered by POC5 domain length. While the relative coverage of the centriole by POC5 is similar between daughter and mother stages, both in control and in *ALMS1^KO^* centrioles. However, the percentage of centriole covered by POC5 is greater in in *ALMS1^KO^*than in control conditions, reflecting that POC5 domain size are similar between stage matched control and *ALMS1^KO^* centrioles while the *ALMS1^KO^* centriole are shorter than control ones. B- Percentage of centriole covered by CEP44 domain. in control conditions, CEP44 domains grows proportionally to the centriole. percentage of daughter *ALMS1^KO^* centriole covered by CEP44 does not differ from that of control. In *ALMS1^KO^* mother centriole however, we detect a small proportional decrease, reflecting the stability of CEP44 domain between daughter and mother stages while centriole size increases. C- Percentage of centriole covered by CCDC77 domain. While the relative coverage of the centriole by CCDC77 increases between daughter and mother stages, there is no difference between stage-matched control and *ALMS1^KO^* centrioles, suggesting that the mechanism controlling CCDC77 domain growth are not impacted by ALMS1 loss.

