## Supplementary figures and images for "ALMS1 contributes to centriole proximal architecture and stability"

### Figure S1

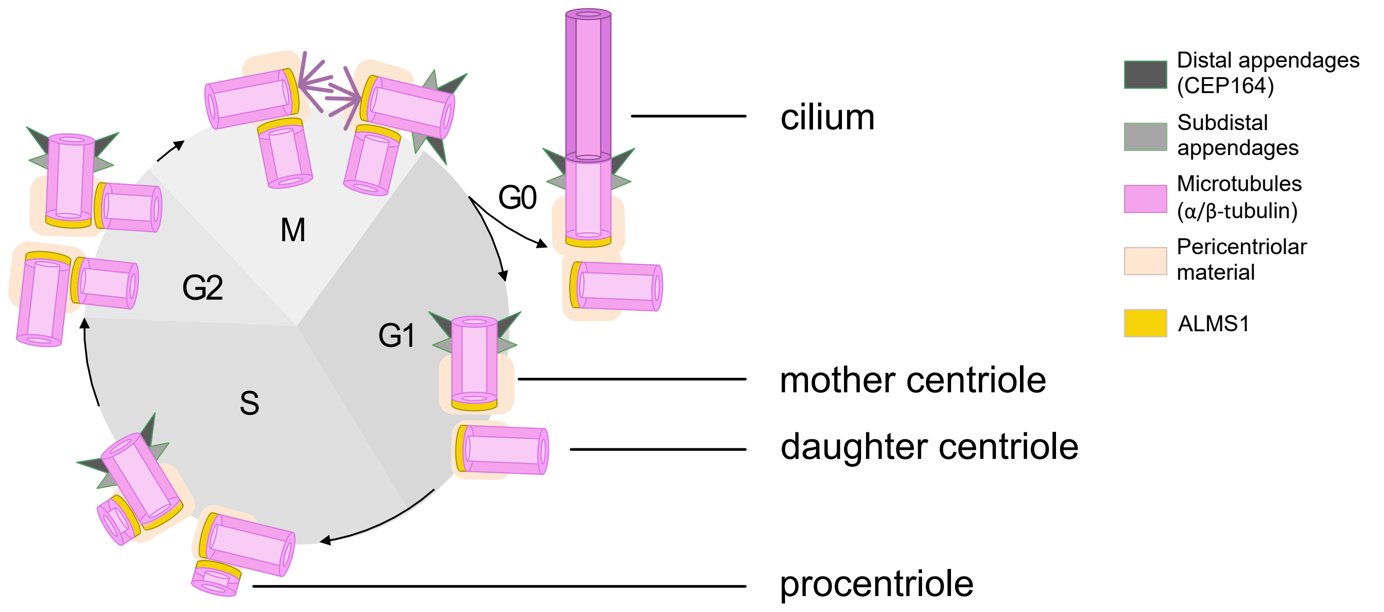

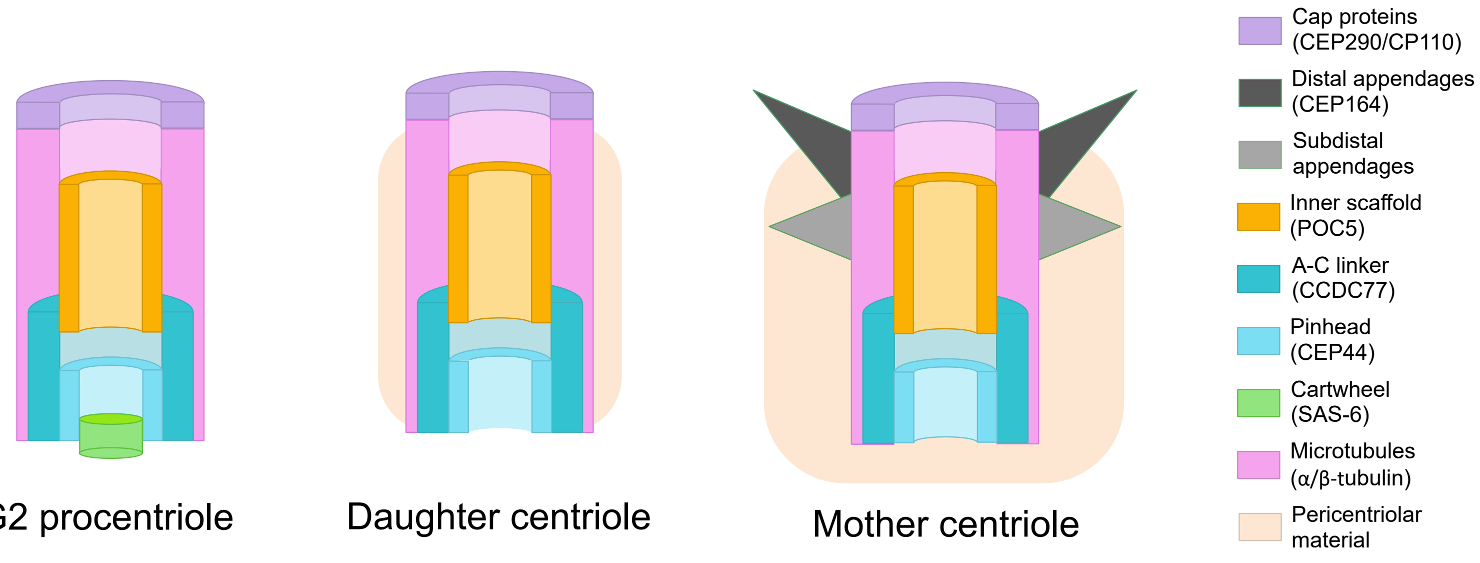

### Figure S2

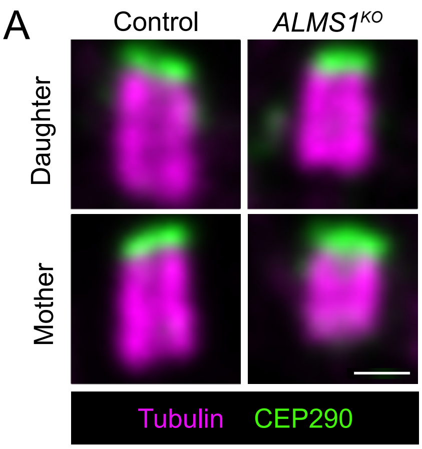


Figure S2

### Figure S3

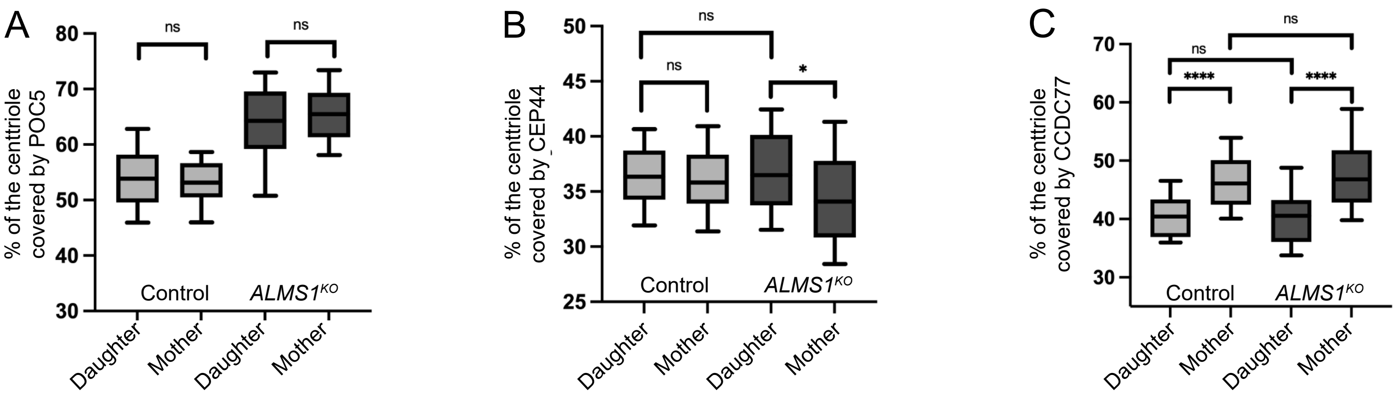
